# Sex-specific microglial responses to glucocerebrosidase inhibition: relevance to GBA1-linked Parkinson disease

**DOI:** 10.1101/2022.12.05.519143

**Authors:** Electra Brunialti, Alessandro Villa, Marco Toffoli, Sara Lucas Del Pozo, Nicoletta Rizzi, Clara Meda, Adriana Maggi, Anthony H. V. Schapira, Paolo Ciana

## Abstract

Microglia are heterogenous cells characterized by distinct populations each contributing to specific biological processes in the nervous system, including neuroprotection. To elucidate the impact of sex-specific microglia heterogenicity to the susceptibility of neuronal stress, we analysed the dynamic changes in shape and motility occurring in primary mouse microglia following pro-inflammatory or neurotoxic insults, thus finding sex-specific responses of microglial subpopulations. Male microglia exhibited a pro-inflammatory phenotype, whereas female microglia showed enhanced neuroprotective capabilities associated with the activation of Nrf2 detoxification pathway in neurons. The sex difference in neuroprotective functions is lost by inhibition of glucocerebrosidase, the product of the GBA1 gene, mutations of which are the major risk factor for Parkinson’s disease (PD). This finding is consistent with the increased risk of PD observed in female carriers of GBA1 mutation, when compared with wild type population, suggesting a role for microglial functionality in the etiopathogenesis of PD-GBA1.

## Introduction

Microglia are resident myeloid cells playing an essential role in the development and homeostasis of the brain, starting from embryonic development and throughout adult life. The physiological function of microglia includes the well-known innate immune response to pathogenic insults ^1^ the sculpting of neuronal termination by pruning synapses ^2^, the engulfment of cellular bodies and debris ^3^ and the synthesis of communication molecules, growth factors and neurotransmitter precursors ^4^, which finally result in a strong influence on synaptic transmission ^5^. The fine tuning of these basic biological processes is ensuring homeostasis and maintains brain tropism, while the presence of dysregulated microglia function is considered a hallmark of neurodegeneration ^6^. The full involvement of microglia in the neurodegenerative processes is still the subject of investigation ^7^, but chronic inflammatory activation may result in neuronal damage ^6^, and abnormal activation of microglia could contribute to the spread of alpha-synuclein and beta-amyloid plaques in the brain of PD and Alzheimer disease (AD) patients ^8,9^. Microglia can exert different functions in the brain by virtue of their marked plasticity, which allows these cells to acquire a wide range of morphological phenotypes, each characterized by different functional properties; these phenotypes can be triggered by specific stimuli, such as pro-and anti-inflammatory cytokines ^10,11^, but are also influenced by the surrounding microenvironment, where the activity of microglia is directed by endocrine ^12^ and paracrine signals ^13^. For these multi-functional abilities, in the different brain areas, heterogenous microglial subpopulations co-exist at the same time ^14,15^. Interestingly, a further level of microglia heterogenicity is due to genetic determinants, including sex-dependent factors which influence both microglia distribution in the central nervous system (CNS) ^16,17^, and some cell-specific morpho-functional properties ^18^, which microglia retain even when transplanted into the brain of the opposite sex ^19^. The differential response to stimulation ^18,20^ of female versus male microglia has been hypothesized to contribute to the sex-dependent bias observed in the prevalence of certain neurological diseases ^12,19,21^, in particular AD and PD for which sex is considered an unmodifiable risk factor ^22,23^. Feminine sex is a risk factor for AD ^23^ and multiple sclerosis ^24^, while male sex is a risk factor for motor neuron disorders ^25^ and PD ^22^. In this context, another clinically relevant (genetic) risk factor for PD is the presence of specific mutations in the GBA1 gene, which have been detected in up to 5-25% of patients ^26,27^. This gene encodes for a lysosomal hydrolase, namely the glucocerebrosidase (GCase): biallelic mutations in GBA1 causes Gaucher Disease (GD) ^28^, while heterozygotic carriers do not develop GD but retain the increased risk to develop PD ^29^. Although most studies previously focused on the functional effects of GBA1 mutations in neurons, our recent investigations revealed that GCase inhibition in microglia is sufficient to impair the physiological ability of microglial to protect neurons against oxidative stress and neurotoxic stimuli ^30^: this acquired microglia phenotype may contribute to the increased risk of neurodegeneration observed in GBA1 carriers.

To investigate the microglial phenotype due to GCase inhibition, in the current study, we developed and applied a non-invasive imaging approach on primary cultures generated by murine models of both sexes. This original methodology allowed us to record in real time the changes of cell morphology induced by specific pharmacological stimuli, with the aim of associating the dynamic variation in cell shape and motility to the biochemical effects induced by the treatments. With this analysis we found that the effects of GCase inhibition in microglia are sex-dependent, thus showing a greater loss of neuroprotective ability of female’s as compared to male’s microglia.

## Results

### Image-based microglia analysis allows detecting functional clusters

To investigate the changes of microglia morphology occurring as a consequence of specific stimuli, we generated an unbiased imaging approach allowing for the dynamic quantification of shape and movement variations of single cells over a fixed period of time. To mimic the physiological microglial environment, we seeded primary adult microglial cells obtained from CX3CR1^+/GFP^ mice, constitutively expressing GFP ^31^, on a layer of neuron-enriched primary culture of cortical cells from syngeneic wild type mice (Supplementary Fig. 1) known to structurally support microglia ^19^. Time-lapse microscopy allowed the recording of morphology and movements of GFP-positive microglia over 2 h; the recorded movies were processed with the ImageJ software ^32^ to obtain morphological and kinetic descriptors for each cell in the acquired field of view (Fig. 1A and Supplementary Movie 1). Briefly, the background was subtracted from the acquired images, which were in turn binarized using a defined threshold that enabled cluster regions of pixels based on similarities threshold to distinguish cell shapes and generate an object for each cell; then the binarized images were processed to remove noise by smooth and despeckle functions of the software to produce sharp objects. A threshold of 130 μm^2^ for the surface size was selected to sort the shapes of cells (microglia) from those originating from cellular debris. The selected shapes were processed to measure two static morphology descriptors: the cell area in square micrometers and the solidity (Fig. 1B) ^33^, the latter defined as the ratio of the area divided by the area of the smallest convex set polygon that contains the cell ^32^, thus resulting in a higher solidity for ameboid rather than for ramified shapes, in a range from 0 to 1 values (Fig. 1B). For each cell, measurements were taken in every frame of the time lapse acquisition; median values of these measurements described the predominant morphology during recording and were used to generate the graphs (Fig. 1C and 1E). Coefficients of variation (CV%) for area and solidity were calculated to obtain numeric descriptors of the dynamic changes occurring during the 2-h measurements (Fig. 1D and 1F). The CV% of the cell area due to size variation was taken as a surrogate marker of cell contractility, while the CV% of the solidity was considered as a measure of the morphological modifications in terms of complexity.

**Figure 1.**
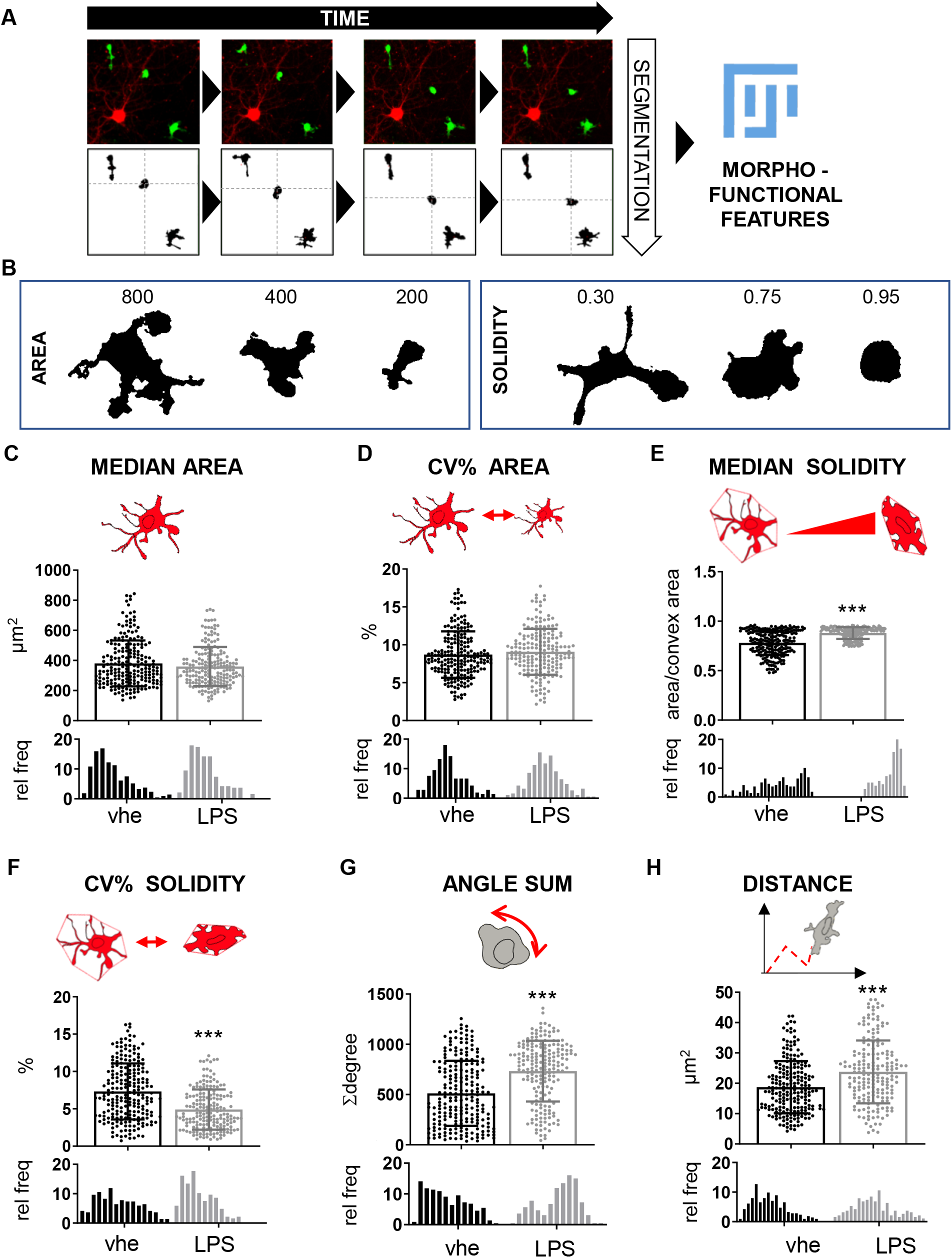
Unbiased morpho-metric method to detect morpho-functional changes. A) Schematic representation of the image-based method for the morpho-functional clustering: GFP-marked microglia dynamics grown on neuronal layer were recorded by time-lapse microscopy, and the acquired images were segmented and processed with Fiji software to obtain shape and dynamic descriptors for each microglial cell during the time. B) Representative images of shape descriptors (area and solidity) used for the analysis. Quantitative single-parameter analysis of male microglia treated with 10 μg/ml lipopolysaccharide (LPS) for a time span of 2 h, starting from 6 h after the treatment; the analysis includes C) median area, D) coefficient of variation (CV%) of the area, E) median solidity, F) CV% of solidity, G) rotation and H) distance covered, the values are presented as mean ±SD (top) and as frequency distribution (bottom) of n=3 independent experiments. The drawings are a schematic representation of the parameter reported in the graph. ***p<0.001 calculated by t-test versus vehicle.

Since microglia are cells able to sense the environment and migrate in response to specific stimuli ^34^ the distance traveled by microglia during the recording time was also considered as a parameter inherently linked to their activity: distance was calculated by tracing the shift in the center of mass of each cell occurring frame by frame, in terms of coordinates (x, y). To define the total covered distance, all shifts were summed and converted into μm values (Fig. 1H). Finally, we measured the number of rotations performed by each cell, another descriptor representing microglial dynamics: in order to calculate this parameter, the ellipse in which the cell can be inscribed was identified and used to calculate the angular displacements frame-by-frame, which were in turn added up to obtain the total rotation of the cells during the recording, expressed as angular degree values (Fig. 1G).

To validate the method we analyzed the descriptor changes associated with a well-characterized microglia polarization, namely the one caused by the potent endotoxin lipopolysaccharide (LPS) ^35^ a strong inducer of a pro-inflammatory microglial phenotype ^36^. Male-derived microglial cells were cultivated on the layer of primary neuron-enriched cultures for 24 h and treated with 10 μg/ml LPS; microglia morphology and motility were analyzed and compared with vehicle-treated cells, by processing videos captured from 6 up to 8 h after the treatment, a time point that is associated with high gain of pro-inflammatory features ^37^. The experiment revealed that the selected descriptors were effective in detecting and describing specific features of microglia induced by LPS (Fig. 1 and Supplementary Movie 2) ^38,39^. In detail, the area of the analyzed cells did not change during the acquisition (Fig. 1C and 1D), but the treatment induced an increase in their solidity of about 13% meaning that when microglia were treated with LPS, their shape became more ameboid (Fig. 1F), while cells maintained their complexity across time, since the variation of solidity (CV% solidity) was higher in vehicle-treated and lower in LPS-treated cells (Fig. 1G). As expected, the cell kinetics was also affected by LPS, showing an increase of about 43% in the number of rotations (Fig. 1G), and of about 27% in the covered distance (Fig. 1H) when compared with vehicle-treated cells.

The results showed that the single-parameter analysis was efficiently identifying phenotypic changes induced by a strong stimulus -as potent as LPS is – occurring in the overall microglial population, but did not provide any detail on the presence of microglia subpopulations with different behavior (Fig. 1 A-H, Supplementary Movie 2). This is particularly important, since microglia shows a peculiar heterogeneity in physiological condition suggesting the existence of various subpopulations reacting differently upon stimulation ^40,41^; we attempted to discriminate these different microglia subpopulations by combining our cellular descriptors in a cluster analysis. We performed a biparametric analysis to test whether we could distinguish the existence of distinct morpho-functional categories: the medians of morpho-dynamic descriptors were used as a cutoff to assign each cell to a descriptive category representative of a value above or under the media. By using the combination of two parameters, cells were clustered into four different subpopulations. As an example, by analyzing solidity and area (Fig. 2A) it was possible to generate four clusters representing microglia subpopulations: Cluster 1) “simple & big”, Cluster 2) “simple & small”, Cluster 3) “complex & big” and Cluster 4) “complex & small”. Cluster 1 is composed of cells that have both area and solidity above the median, in contrast, the cells with area and solidity under the media fall in Cluster 4; cells with the bigger area and low solidity fall in Cluster 3, and the cell small and simple in Cluster 2. The categories generated for each parameter are reported in Table 1 and Supplementary Table 1. The application of this approach to the data obtained with the LPS experiment, in keeping with previous reports ^19,38,39,42^, revealed that after treatment a population of cells characterized by a “complex & small” shape mostly disappeared, while an increase in the subpopulation of “simple & small” cells was observed (Fig. 2A) and the subpopulation of “simple & big” cells, which was under-represented in the vehicle-treated samples, became prominent after LPS treatment (Fig. 2A). The cluster analysis was applied to identify different morpho-functional subpopulations and was reported in specific histograms (Fig. 2B) demonstrating how the different subpopulations were affected by the LPS treatment. The graphs show that LPS treatment increased the subpopulation of “small & motile”, “big & motile”, “steady & motile”, “contractile & simple”, “simple & small”, “simple & big”, “simple & motile” and “rotant & motile” (Fig. 2B). Interestingly, the method was able also to detect that some categories of cells did not respond to LPS and their subpopulation remained unaffected after the treatment, e.g. “steady & static”, “variable & motile”, “complex & motile”, “simple & static” (Fig. 2B). These results demonstrated that the dynamic morpho-functional analysis allowed to discriminate microglial subpopulations differentially responding to specific stimulations.

**Table 1:** For each parameter, using as a cut-off the median we generated two groups named as reported in the table. Therefore a microglial cell is included in one or the other group according to the value of the descriptor if it is bigger or lower respect to the median value.

| PARAMETER | GROUPS GENERATED |  |
| --- | --- | --- |
|  | UNDER THE MEDIAN | OVER THE MEDIAN |
| MEDIAN AREA | BIG | SMALL |
| CV% AREA | INACTIVE | CONTRACTILE |
| MEDIAN SOLITIDY | COMPLEX | SIMPLE |
| CV% SOLIDITY | STEADY | VARIABLE |
| MEDIAN MOTILITY | STATIC | MOTILE |
| MEDIAN ROTATION | STATIONARY | ROTANT |

**Figure 2.**
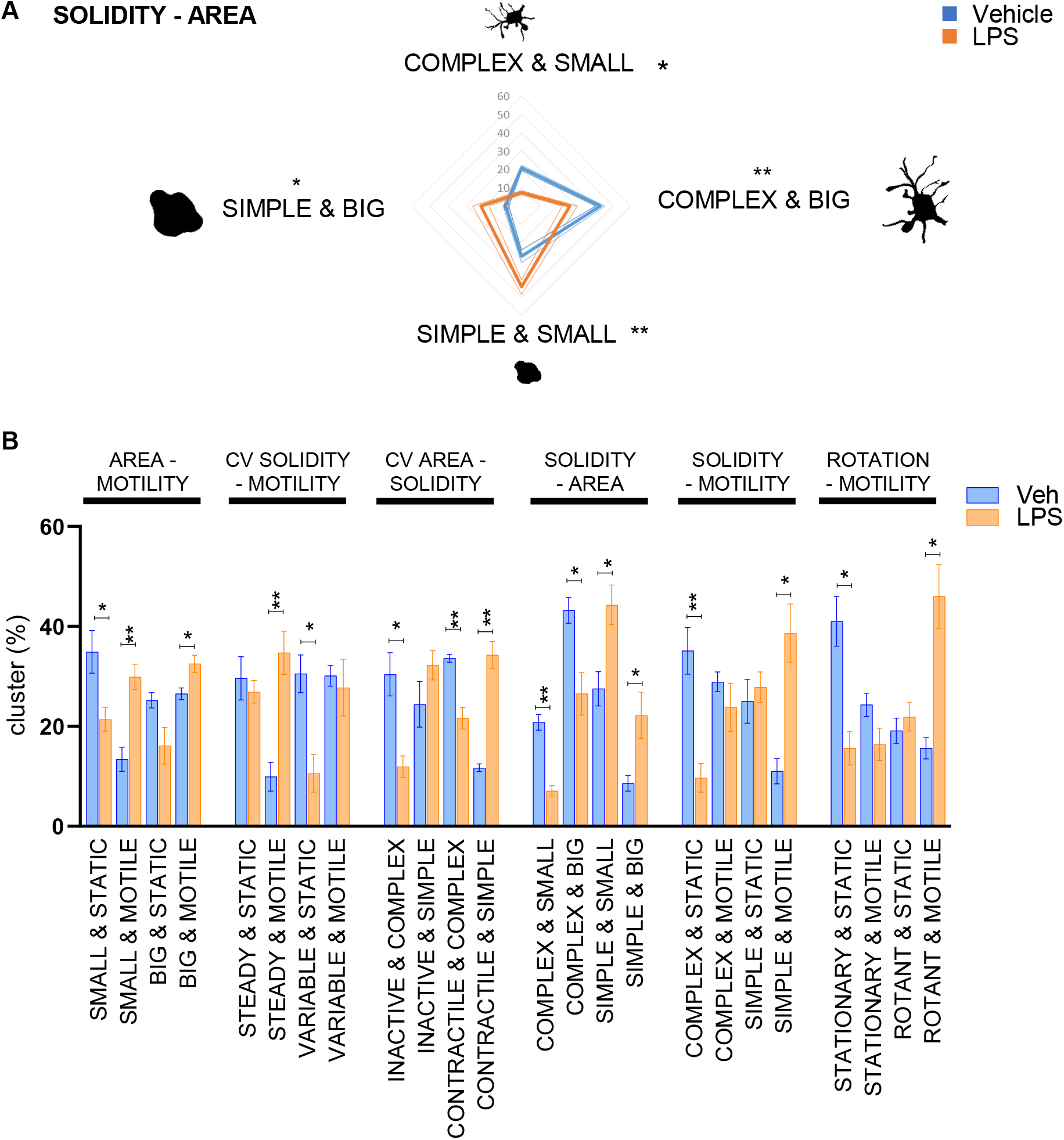
Morpho-functional biparametric analysis of LPS treated microglia. Analysis was carried out in the time interval 6-8 h after treating cells with 10 μg/ml LPS. A) Biparametric analysis of microglial solidity and area allows to cluster the cells into four categories: “complex & big”, “complex & small”, “ simple & small”, “simple & big”; the percentage of each population is reported as radar graph relative to 3 independent experiments. Drawings represent the morphology representative of the category. *p<0.05; **p<0.01; ***p<0.001 calculated by t-test versus vehicle. B) biparametric analysis represented as histogram of the median ± SEM of the sub-population percentage obtained in 3 different experiments, the two parameters obtained to generate the clusters are reported on the top of the graph; *p<0.05, **p<0.01,***p<0.001 calculated by t-test versus vehicle.

### Male- and female microglia show different morpho-functional phenotypes

Once validated, the morphometric approach was used to test whether sex-differences could be detected in the dynamic behavior of microglia. To this end, primary brain microglia cells from male or female mice were isolated from adult CX3CR1^+/GFP^ and seeded on neuron-enriched primary cortical cells from syngeneic wild type mice (mixed population of male and female mice). 24 h after seeding microglia dynamics were recorded for 2 h to identify possible sex-related differences in unstimulated conditions. Indeed, different subpopulations were present in female and male microglia: when compared to male, female microglia showed cluster subpopulations of “big & static”, “variable & static”, “inactive & complex”, “complex & small”, “complex & static”, and “rotant & static” (Fig. 3). These data suggested that female microglia in physiological conditions are enriched in subpopulations characterized by complex and static cells, possibly interacting with the surrounding micro-environment, with a less pro-inflammatory profile: this is in accordance with previous reports indicating that, in female mice microglia are more dedicated to the maintenance of brain homeostasis, while male microglia are more inclined to perform defensive tasks ^19^.

**Figure 3.**
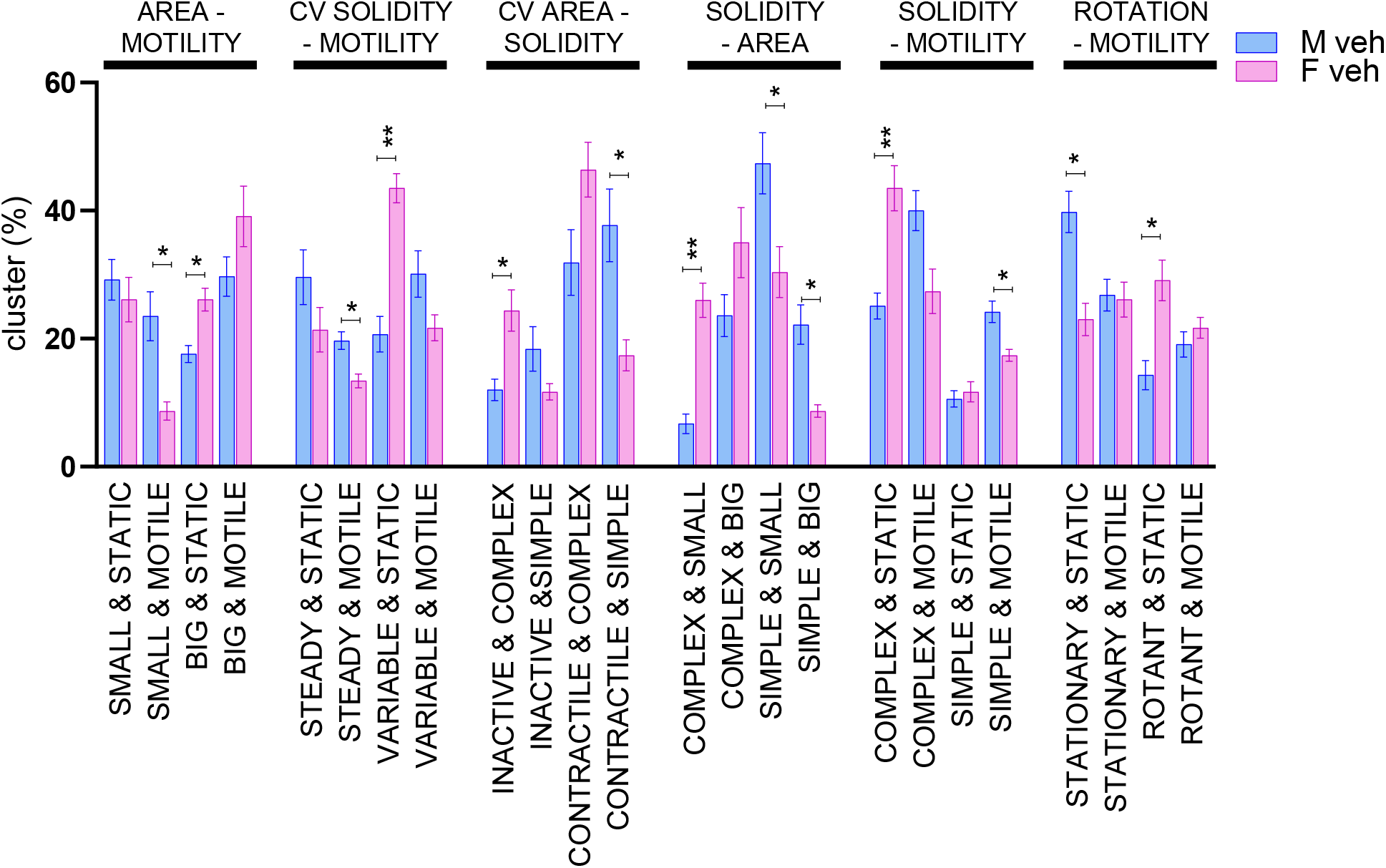
Morpho-functional biparametric analysis of male and female microglia. Biparametric analysis is represented as histogram of the median ± SEM of the sub-population percentage obtained in 3 different experiments, the two parameters obtained to generate the clusters are reported on the top of the graph; *p<0.05, **p<0.01 calculated by t-test versus vehicle.

### Chemical inhibition of β-glucocerebrosidase (GCase) exerts a differential effect in male and female microglia

We previously demonstrated that the pharmacological inhibition of microglial GCase with conduritol-B-epoxide (CBE) interferes with the neuroprotective function of microglia ^30^. To characterize the microglia morphology in response to GCase inhibition, we carried out the morpho-functional analysis after treating cocultures with 200 μM CBE: this concentration was selected in order to ensure a almost total (−98% activity) inhibition of GCase activity sufficient to selectively interfere with the microglia neuroprotective functions ^30^, while with negligible effects on the activity of additional glycosidase targets ^43,44^. The dynamic changes of microglial morphology were recorded at early time points (48 to 50 h after treatment), a time window in which neuronal and microglia mortality due to CBE were virtually absent ^30^. The morphometric analysis revealed that GCase inhibition of male microglia changed the phenotype of specific sub-populations, increasing cells characterized by a static and less contractile phenotype (Fig. 4A). We observed increases of cell populations with “big & static”, “steady & static”, “simple & big “, “simple & static”, “stationary & static” while a decrease in the sub-populations of cells “big & motile”, “variable & motile”, “contractile & complex”, “complex & motile” and “stationary & motile” (Fig. 4A).

**Figure 4.**
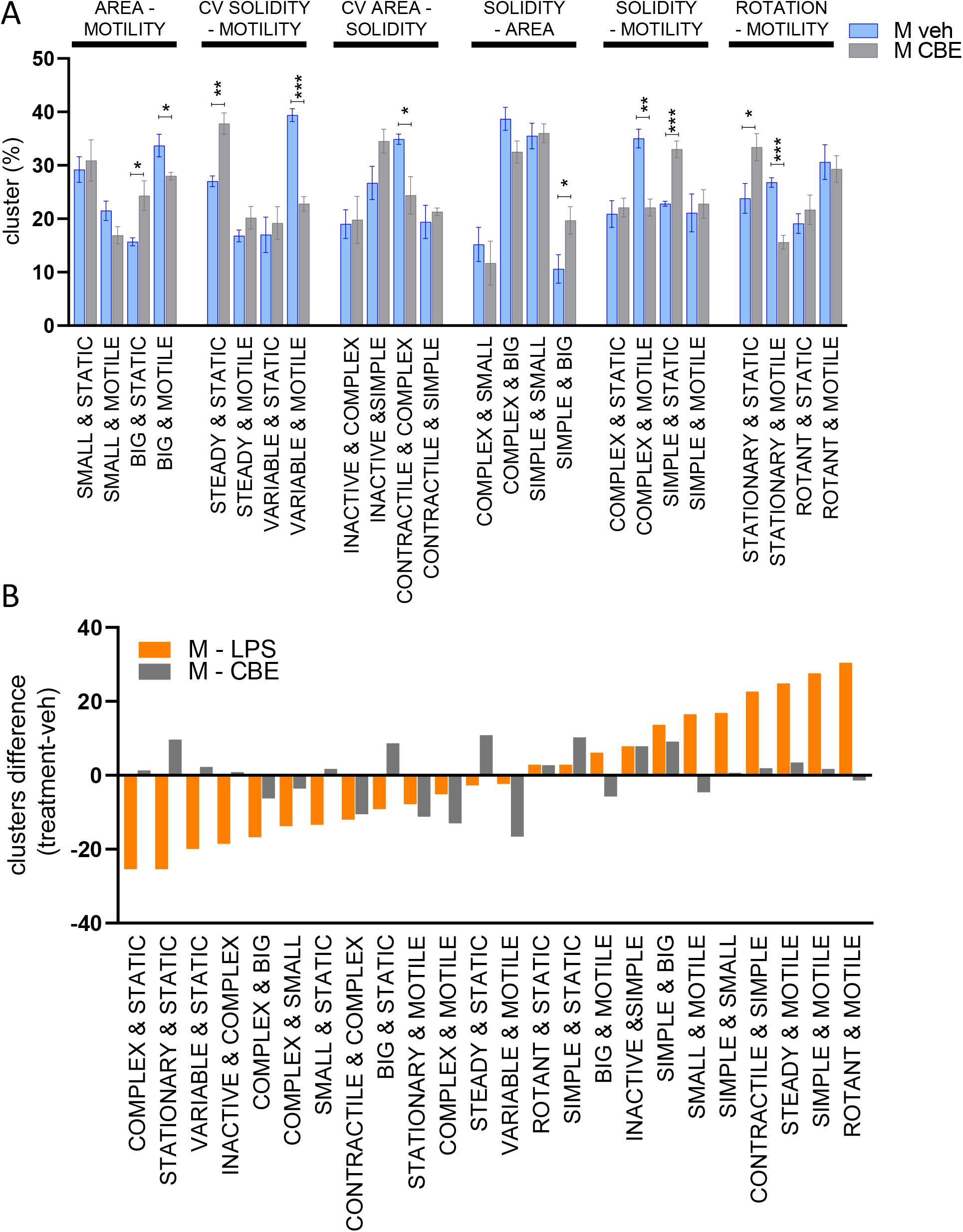
Morpho-functional biparametric analysis of CBE-treated male microglia. Analysis was carried out in the time interval 48-50 h after treating cells with 200 μM CBE. A) Biparametric analysis is represented as a histogram of the median ± SEM of the sub-population percentage obtained in 4 different experiments, the two parameters obtained to generate the clusters are reported on the top of the graph; *p<0.05, **p<0.01, ***p<0.001 calculated by t-test versus vehicle. B) Variation in subpopulation composition for LPS (Figure 2B) and CBE (A) treatment versus the vehicle-treated cells.

Overall, the phenotype observed was characterized by a large and static morphology (Supplementary Movie 3), inclusive of large and simple shapes associated with a low motility. Since it has been reported that microglia can acquire a pro-inflammatory phenotype after long-term GCase inhibition ^45,46^, we compared the phenotype triggered by LPS (Fig. 2B) with the short-term treatment with CBE (48-50 h). Surprisingly, the morpho-functional analysis of male microglia treated with LPS or CBE (Fig. 4B) demonstrated opposite effects by increasing (LPS) or decreasing (CBE) the motility in most subpopulations.

A sex-difference in microglial reactivity has been previously described ^19^, so we investigated if the CBE treatment differentially affected the microglia phenotype obtained from female or male mice. Thus, we treated cocultures of female microglia with CBE and recorded the effects at the same time points (48 to 50 h after treatment) as the previous experiment. As reported in Fig. 5A, GCase inhibition induced a radical shift in female microglia morpho-functionality, leading to an increased representation of subpopulations of “small & motile”, “steady & static”, “contractile & complex”, “simple & big”, “simple & static” and a decrease in the subpopulations of “big & static”, “steady & motile”, “variable & motile”, “inactive & complex”, “complex & small”, “complex & static”, “complex & motile” and “rotant & motile” (Fig. 5A) cells. Comparison of the results obtained with female (Fig. 5A) and male microglia (Fig. 4A) after CBE treatment revealed that some subpopulations were differentially enriched, and that CBE treatment induced more marked morpho-functional changes in female microglia (Fig. 5B).

**Figure 5.**
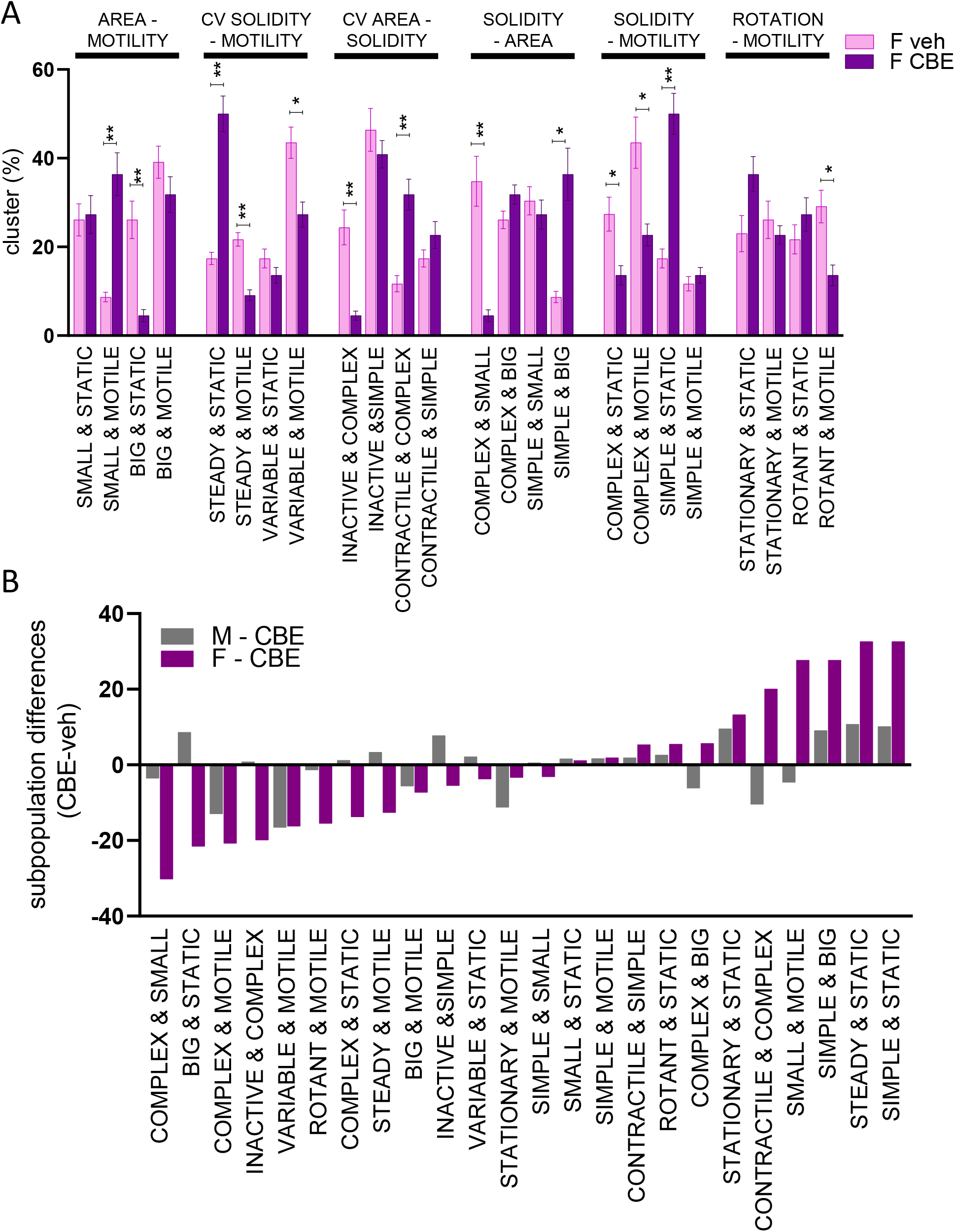
Morpho-functional biparametric analysis of CBE-treated female microglia. Analysis was carried out in the time interval 48-50 h after treating cells with 200 μM CBE. A) Biparametric analysis represented as a histogram of the median ± SEM of the sub-population percentage obtained in 3 different experiments, the two parameters obtained to generate the clusters are reported on the top of the graph; *p<0.05, **p<0.01 calculated by t-test versus vehicle. B) Variation in subpopulation composition for CBE treated male (Fig. 4A) and CBE treated female (A) microglia versus the vehicle treated cells.

Based on these results, we decided to test whether this more pronounced effect of CBE on female microglia also reflected alterations in their neuroprotective functions; indeed, we previously demonstrated that microglia are able to increase neuronal NRF2 transcriptional activity that protect neurons from neurotoxin effects, a mechanism which is impaired by GCase inhibition ^30^. We purified microglia from groups of female and male wild type mice treated with vehicle or 100 mg/kg CBE for 3 days to inhibit microglial GCase ^30^, the purified microglia was seeded over a neuronal cell layer derived from ARE-luc2 mice (Fig. 6A), transgenic animals in which a luciferase reporter is expressed under the control of the Nrf2 transcription factor ^47,48^. This system allowed us to measure the ability of microglia to increase the neuronal Nrf2 activity simply by measuring the luciferase activity in the coculture. Interestingly, female microglia extracted from vehicle-treated mice revealed a more prominent effect in inducing Nrf2 response when compared to male microglia. The effect of CBE treatment, that is expected to reduce neuronal to microglia Nrf2 response ^30^, was sufficient to blunt the differences observed between male and female microglia obtained from vehicle-treated mice (Fig. 6B), thus suppressing the neuroprotective action exerted by microglia independently from the sex of origin.

**Figure 6.**
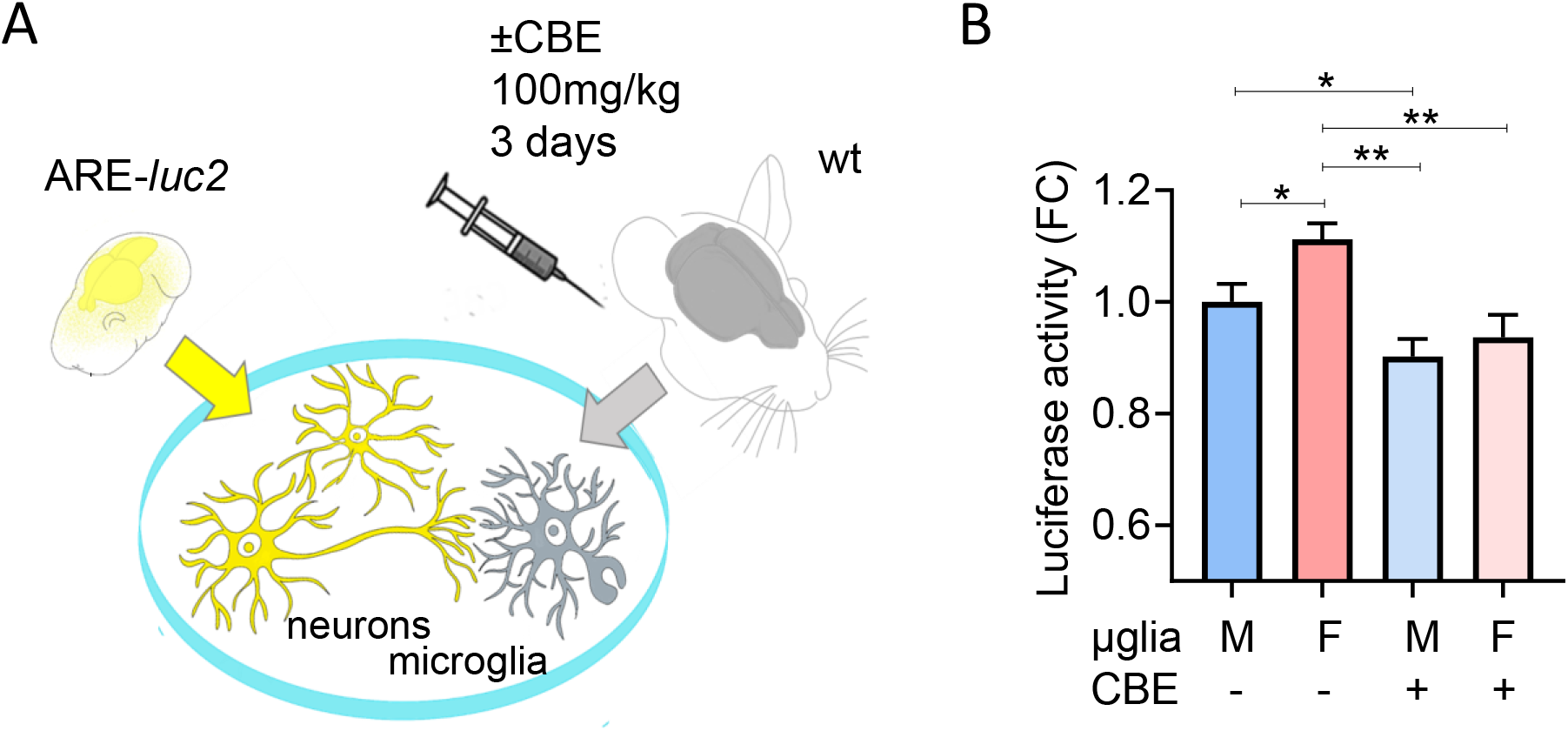
CBE treatment reduces the neuronal NRF2 response induced by microglia. A) Scheme of the experiments reported in B, aimed at testing the effect of primary microglia (μglia) extracted from male or female mice treated with 100 mg/kg/die CBE for 3 days. B) Luciferase activity measured in protein extracts derived from ARE-luc2 neurons cultured with microglia derived from CBE-or vehicle-treated mice (A). Data are presented as mean ±SEM of n=7 independent samples. *p<0.05, **p<0.01 calculated by unpaired t-test vs the corresponding sample.

Based on these data, which suggest that a reduction in GCase activity decreases the protective microglial response in female mice, we hypothesized that the normal male predominance seen in PD patient populations would be abolished in PD subjects with GBA1 variants.

We analyzed the AMP-PD database that includes a total of 3497 individuals (Fig. 7). Of these, 1971 (56.4%) were males and 1526 (43.6%) were females. For idiopathic PD cases alone, there were 1236 males (63.4%) and 715 females (36.6%). In the *GBA*-PD group there were 163 males (57.4%) and 121 females (42.6%). Statistical analysis showed that the male predominance in idiopathic PD is lost in *GBA1*-associated PD, although this just fails significance at p=0.0525.

**Figure 7.**
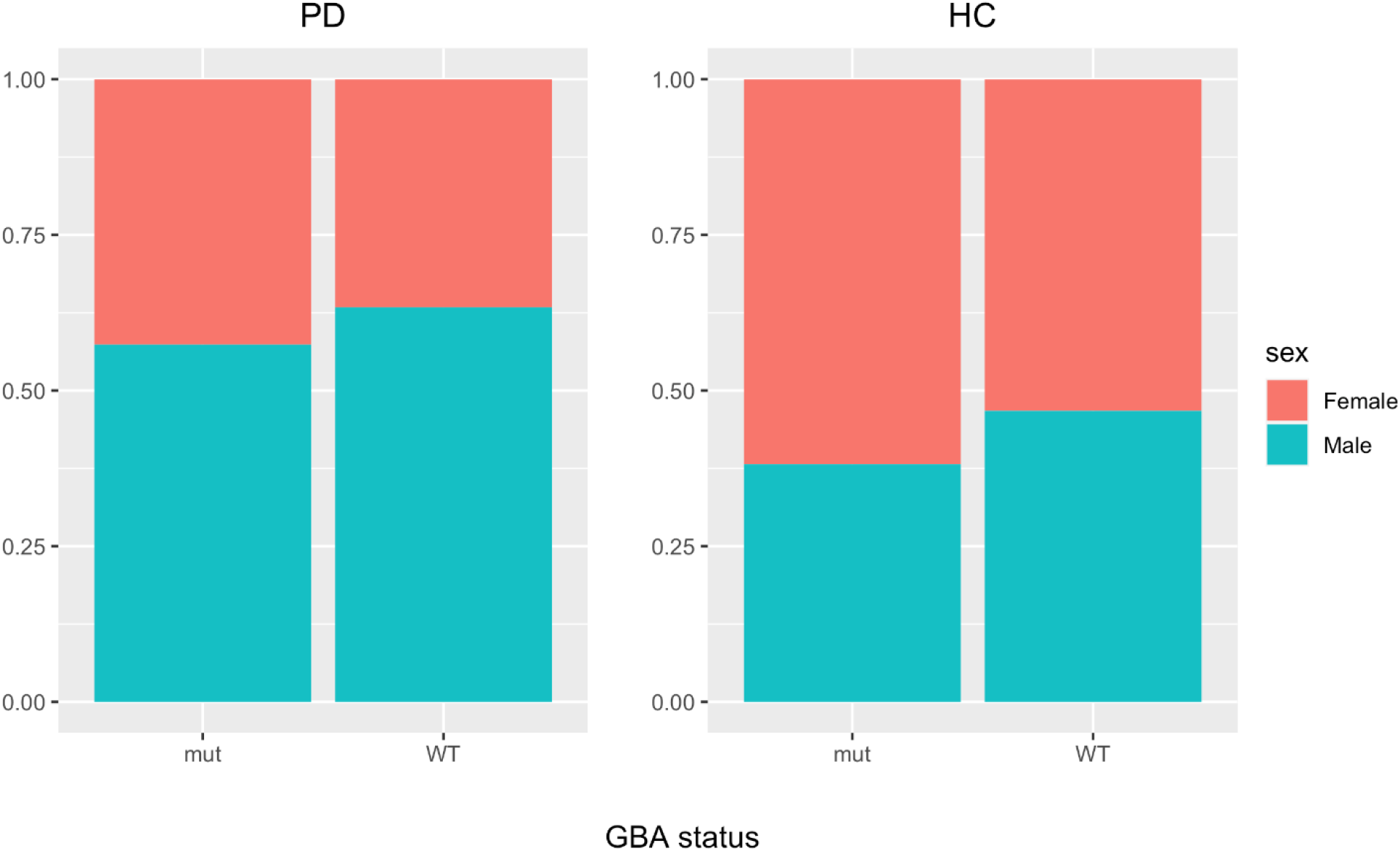
Different sex distribution in idiopathic-PD and GBA-PD patients. In the plot are summarized the ratio of females and males in PD and healthy control (HC) populations; mut = mutant.

## Discussion

Microglial cells are characterized by the presence of different subpopulations, which differ in abundance and morphology, and are characterized by distinct genetic programs, protein expression patterns, and ability to respond to environmental stimuli ^14,15^. The distribution of these subpopulations follows a spatial-temporal definite pattern: indeed, specific phenotypes can be detected at different evolutionary stages, but they can also coexist simultaneously in brain parenchyma of adult animals ^40,49^. The morphology of microglia is indicative of their functional status ^10,50^, thus analysis of microglial shape can anticipate information about the biochemical pathways triggered in these cells by pathophysiological processes ^51^. Standard morphological analysis based on immunocytochemistry images provides snapshots of cell shape and offers a static view of the cell population ^33^ but it does not detect dynamic changes, such as the variation in cell protrusions or changes in migration, features that are certainly key components of microglial biology and allow better deciphering their behavior ^52^. In our study, we added the temporal dimension to standard shape analysis by applying time-lapse fluorescence microscopy to our *in vitro* model of the multicellular condition of the brain, encompassing a co-culture of GFP-expressing primary microglia and primary cortical cells enriched in neurons. In this context, we applied an unbiased imaging-based analysis for each microglia cell of the investigated population, and for each frame of the recorded movies, we measured standard morphological cues that included cell dimension and complexity, together with novel dynamic descriptors able to describe time-dependent changes in microglia motility, contractility, rotation, and complexity. The method was effective for detecting changes occurring in pro-inflammatory microglia, which have been previously described ^38,53^ and include changes in the cellular shape towards the amoeboid morphology, and a general increase in the motility (Fig. 1 and 2).

The first set of experiments was designed to validate the method, and the results were in line with prior knowledge, but at the same time revealed information on the response of primary microglia to pro-inflammatory stimuli by disclosing resilient subpopulations of cells that did not undergo substantial changes after stimulation. These subpopulations show phenotypes that – following the classification generated by our protocol -are defined by the descriptors “complex & motile”, “simple & static”, “stationary & motile” and “rotant & static”, and display a non-responsive phenotype against LPS stimuli (Fig. 2B). Thus, our morpho-functional analysis provided a direct demonstration that adult microglia exist in different subtypes, each characterized by peculiar shapes, and possibly by different gene expression profiles and function ^40,51^, and that these subtypes can differently respond to stimulations.

Once validated, we have applied the morpho-functional analysis to evaluate sex-related differences in the composition of microglia subpopulations. In previous studies, biochemical, morphological, and functional data recognized sex-related differences in microglia revealing that male microglia have a constitutive mild pro-inflammatory phenotype, while female microglia are reminiscent of surveilling microglia, that for definition are stationary and ramified cells sensing the environment ^19,54,55^; these differences were shown to be genetically determined, independent of hormonal status and of the microenvironment, indeed are maintained also when microglia are maintained in culture, when cross-transplanted in a brain of opposite sexes and when microglia were extracted from brains of ovariectomized females, where the hormonal environment was similar to male mice ^19^. It is likely that these sex-related differences in microglia are contributing to the differential sex-specific susceptibility and severity of some neurological diseases ^12,56,57^.

In our morpho-functional analysis, male microglia, when compared to female cells were enriched in sub-populations defined by the descriptors “small & motile”, “simple & motile”, “contractile & simple”, “steady & motile”, “small & simple”, “simple & big” (Fig. 3), which are increased when cultures are treated with LPS (Fig. 2B), thus supporting the notion that male microglia show a higher tendency to acquire a pro-inflammatory phenotype than female microglia. In contrast, in female microglia we found a marked presence of subpopulations defined by the descriptors “big & static”, “variable & static”, “inactive & complex”, “complex & small”, “complex & static” (Fig. 3), categories that are decreased after LPS treatment (Fig. 2B) supporting the hypothesis that female cells show a less pro-inflamed phenotype, with a profile reminiscent of surveilling microglia ^10,19^.

Do these differences influence the development and progression of a neurodegenerative process? Previous data from our lab showed that microglia may contribute to brain neuroprotection by inducing the Nrf2 pathway in neurons through direct contact between microglial cells and neurons. This Nrf2-activation is reduced when microglial GCase is pharmacologically inhibited, an effect that renders dopaminergic neurons more sensitive to neurotoxic stimulations ^30^. On the basis of these data, we hypothesized that the reduced microglial neuroprotective functions might contribute to the observed increased risk of PD in carriers of GBA1 mutations and prompted us to analyze the microglial morpho-functionality after GCase inhibition. Interestingly, the morpho-functional analysis on static descriptors demonstrated that inhibition of GCase enriched the microglial population with cells characterized by an ameboid-like (“simple & big”) morphology like those observed with LPS (Fig. 4) and typical of the pro-inflammatory activation. Similar peculiar microglial morphologies with bigger soma and less protrusion have been detected in murine and vertebrate GCase deficient models induced by genetic modification ^58,59^ and in brain areas (such as *substantia nigra*) of neuropathic GD patients ^60,61^, and were often associated with a pro-inflammatory phenotype. However, with our morpho-metric analysis, the use of dynamic descriptors clearly distinguished the effects of LPS and CBE treatments on microglia, showing that male CBE-treated cells were static and less contractile, a phenotype markedly different from the pro-inflammatory phenotype (Fig. 2B and 4B), and characteristic of inactive microglia. This is in line with our previous expression data showing that no pro-inflammatory genes were induced by the CBE treatment in immortalized microglia ^30^. The more stationary phenotype and the decreased number of protrusions suggest that CBE-treated microglia display a reduced contact surface with the neuron membranes, a condition likely contributing to the decreased Nrf2 expression in neurons ^30^. The reduction of Nrf2 levels might increase the risk of neurodegeneration especially in neurons of the *substantia nigra* that are frequently exposed to oxidative stress due to dopamine metabolism ^62^. In the case of GBA1 mutations, the microglial GCase inactivation is constitutive and over time could promote pathways leading to promotes neurodegeneration.

Interestingly, female microglia seem to be more affected by GCase impairment, indeed the morpho-functional phenotype is more divergent from the vehicle when compared to male microglia (Fig. 5): the CBE effect on female microglia increases the subpopulations characterized by a less active behavior (less ramified shape and static) to a greater extent compared to male microglial cells.

Moreover, female microglia also displayed a divergent response as compared to male microglia, indeed CBE treatment increased the subpopulations defined by the descriptors as “small & motile” and “complex & contractile”; to our knowledge, this is the first description that the inhibition of GCase is able to have a different effect on the morphology and motility of male and female microglia.

Intriguingly, the effect of GCase inhibition is more penetrant in female microglia, reducing the superior ability of female microglia to induce Nrf2 in neurons to the same extent found in male microglia (Fig. 6). This dramatic change in female microglia function is likely diminishing the greater neuroprotective ability of female microglia, rendering them comparable to male microglia. These sex-related morpho-functional differences may have functional consequences: it is tempting to speculate that the increased neuroprotective ability of female microglia could contribute to the 1.5-2 fold reduced risk of developing idiopathic PD observed in female individuals, a sex-bias which appears to be reduced in GBA1-PD patients (Fig. 7) ^63–67^. Indeed, the majority of studies report higher female prevalence in GBA-PD, or do not observe sex-related differences ^64–67^ suggesting that the protective effect associated with female sex is indeed blunted by GBA mutations ^65^, although a firm explanation of this difference with idiopathic PD have not been reported. Our data suggest that the differential effects of these mutations on the microglial phenotype might contribute to the differences observed between idiopathic and GBA-PD in terms of loss of male predominance.

In conclusion we report a novel methodological approach toward the identification of dysfunctions of microglia in models of neurological diseases. The morpho-functional method was revealed to be sufficiently sensitive to recognize phenotypic differences in unstimulated microglia derived from the brain of male or female animals; moreover, the technique demonstrated the existence of discrete subpopulations of microglia, each characterized by specific morphological descriptors, indicative of different specific phenotypes and a differential response to specific stimuli. This novel perspective provides insight into the microglial heterogeneous behaviors that might underlie pathological stimuli in different CNS regions or in the function of sex and age ^68^. Indeed, the identified morpho-functional parameters allowed us to describe the morphological changes induced not only by a well-known pro-inflammatory agent (LPS), but also by the CBE model of reduced GCase activity and GBA1-PD. Our data, for the first time demonstrates that GCase inhibition triggers a specific microglial morpho-functional phenotype associated with a reduced ability of microglia to perform neuroprotective functions, with more dramatic consequences for microglia isolated from female animals: this finding might contribute to the understanding of the sex-related differences clinically observed in idiopathic PD.

## Methods

### Cell cultures

Primary neurons were derived from the cerebral cortex of p0-p1 mice following standard operational procedure using the neural tissue dissociation kit-postnatal neurons (Cat. 130-094-802, Miltenyi Biotec) as previously described ^30^. In brief, the brain cortices from six mice were pooled as a single experimental group and subjected to enzymatic and mechanical dissociation, then 150,000 primary neuronal cells were seeded for each well of a poly-L-ornithine coated 24-well plate, replacing half of the medium volume every 2 or 3 days. At day ten, 37,500 primary microglia cells isolated from the whole brain of adult mice (age 3-6 months) were seeded on neuron layer; briefly, the brains from two mice were pooled and subjected to enzymatic and mechanical dissociation and microglia were purified using a magnetic column and CD11b coated microbeads (Cat. 130-093-634, Miltenyi Biotec) ^19^. Neuronal and microglial cultures were grown in Neurobasal A medium (Cat. 10888-022, LifeTechnologies) containing 1% streptomycin–penicillin, 1 % GlutaMAX, 2% B-27 Supplement (Cat. 17504-044; Gibco), 10 mM HEPES (Cat. H0887, Merk), in a humidified 5% CO2-95% air atmosphere at 37 °C.

### Cell treatments

For lipopolysaccharide (LPS) experiments, cultures were treated with a final concentration of 10 μg/ml LPS O111:B4, (Cat. L2630, Merk) for 6 h or vehicle (water); for the CBE experiments, cultures were treated with a final concentration of 200 μM CBE (Cat. 234599, Merk) or vehicle (water) for 48 h and then subjected to timelapse microscopy.

### Fluorescent image acquisition and processing

Time-lapse sessions were performed on live microglia for 20 random fields per condition using an Axiovert 200M microscope with dedicated software (AxioVision Rel 4.9, Zeiss, RRID:SCR_002677, https://www.micro-shop.zeiss.com/it/ch/system/software+axiovision-axiovision+programma-axiovision+software/10221) at ×20 magnification; the recording was performed for 2 h taking a picture every 5 min. An algorithm was generated to segment and analyze GFP positive cells (namely microglia) exploiting Fiji software (ImageJ, NIH, version 2.0.0, RRID:SCR_002285, http://fiji.sc). The background was subtracted and set constant across the experimental groups; the class of pixels with a value over the defined threshold (foreground) that corresponds to green fluorescent objects have been subject to the despeckle and smoothing function. Then the objects with an area greater or equal to 130 μm^2^, were subjected to the “analyze particle” function to calculate the “Area”, “Center of mass”, “Shape descriptors” and “Feret’s diameter” for each object in each frame. The area was converted from pixel to the surface in μm^2^; the coordinates of the center of mass of each object were used to calculate the distance covered by the cell during the time-lapses. Among the “Shape descriptors”, we operated with the solidity, a value that corresponds to the area/convex area of the object; between the “Feret’s diameter” values we used the “Feret Angle” to calculate the number of rotations of each object during the recording. An math operation was used to perform the clustering analysis: in brief, for each parameter obtained from the analysis the values of the vehicle and treated cells were used to identify the median parameter for the experiment, this median was used as a threshold to cluster the cells in two groups (over or under the median); the combination of two parameters was used to generate four different clusters (Table 1 and Supplementary Table 1).

### Animals and treatments

The animals were fed ad libitum and housed in individually ventilated plastic cages within a temperature range of 22–25 °C under a relative humidity of 50% ± 10% and an automatic cycle of 12 hours light/dark. C57BL/6 and CX3CR1^+/^GFP mice were supplied by Charles River (Charles River Laboratories MGI Cat 2159769, RRID:MGI:2159769 and MGI:J:84544), ARE-luc2 mice were generated in our laboratory ^47^. For pharmacological treatments, mice (15–30 weeks old) were administered 100 mg/kg/day CBE or vehicle (PBS) via i.p. injection for 3 days before the purification of microglia.

### Luciferase enzymatic assay

Luciferase assays were performed as illustrated previously ^48^ in brief, microglia-neuron cultures were lysed with luciferase cell culture lysis reagent (Cat. E1531, Promega), and the protein concentration was determined with a Bradford assay^69^. The luciferase activity assay was carried out in luciferase assay buffer by measuring luminescence emission with a luminometer (Veritas, Turner) for 10 s to obtain the relative luminescence units (RLU).

### Clinical data

Clinical data were obtained from the Accelerating Medicines Partnership Parkinson’s Disease (AMP-PD) knowledge portal, downloaded on the 28th May 2020 (release 15^th^ October 2019). Correct GBA sequencing was obtained using Gauchian (Version 1.0.2, https://github.com/Illumina/Gauchian), a software described in a previous publication ^70^.

Participants with the following tags were included: “PD”, “Genetic Registry PD”, “Genetic Cohort PD”, “Genetic Registry Unaffected”, “Genetic Cohort Unaffected”, “Healthy Control”. Participants marked as “Prodromal” and “SWEDD” (scans without evidence for dopaminergic deficit) were excluded from the analysis.

### Statistical analysis

For the cellular experiment statistical analyses was performed employing Prism 7 (Version 7.00, GraphPad Software Inc., RRID:SCR_002798, http://www.graphpad.com), multiple t-test versus vehicle were used to determine if there were significant differences in means and a *p-*value lower than 0.05 was considered to indicate statistical significance. For the clinical data statistical analysis was carried out using R (version 4.2.1, RRID:SCR_001905, http://www.r-project.org). Pearson’s Chi-squared test was used to compare sex differences between carriers and non-carriers of GBA variants.

### Data availability

The data that support the findings of this study are available at DOI 10.5281/zenodo.7360295. The following protocols are available at protocols.io: Luciferase Activity Assay (DOI: dx.doi.org/10.17504/protocols.io.j8nlkw8bxl5r/v1); Cell treatments (DOI: dx.doi.org/10.17504/protocols.io.ewov1o98plr2/v1); Fluorescent image acquisition and processing (DOI: dx.doi.org/10.17504/protocols.io.ewov1o98plr2/v1). Other information is available from the senior author upon reasonable request.

## Acknowledgments

The authors are indebted to Finlombarda and the TOPsrl research team for generating the ARE-luc2 reporter mice.

## Author Contributions

Conceptualization, methodology, validation and formal analysis: P.C., E.B., A.V., and NR; investigation: E.B., MT, SLDP, CM and A.V.; Funding acquisition: P.C and AS.; writing of original draft: P.C., E.B., and A.V.; review and editing of the manuscript: P.C., A.S. All authors read and approved the final manuscript.

## Competing interests

The authors declare no competing interests.

## Additional information

### Supplementary information

The online version contains supplementary material available at…

### Funding Information

The work was supported by the EU Joint Programme -Neurodegenerative Disease Research (JPND) project (GBA-PaCTS, 01ED2005B and GBA-PARK n. 212) (to P.C. and A.S). AS is supported by a grant from ASAP. This research was funded in whole or in part by Aligning Science Across Parkinson’s [ASAP-000420] through the Michael J. Fox Foundation for Parkinson’s Research (MJFF). For the purpose of open access, the author has applied a CC BY public copyright license to all Author Accepted Manuscripts arising from this submission.

### Institutional Review Board Statement

All animal experimentation was carried out in accordance with the Animal Research: Reporting of In Vivo Experiments (ARRIVE) guidelines and the European Guidelines for Animal Care. All animal experiments were approved by the Italian Ministry of Research and University.

“Data used in the preparation of this article were obtained from the Accelerating Medicine Partnership® (AMP®) Parkinson’s Disease (AMP PD) Knowledge Platform. For up-to-date information on the study, visit https://www.amp-pd.org.

“The AMP® PD program is a public-private partnership managed by the Foundation for the National Institutes of Health and funded by the National Institute of Neurological Disorders and Stroke (NINDS) in partnership with the Aligning Science Across Parkinson’s (ASAP) initiative; Celgene Corporation, a subsidiary of Bristol-Myers Squibb Company; GlaxoSmithKline plc (GSK); The Michael J. Fox Foundation for Parkinson’s Research ; Pfizer Inc.; Sanofi US Services Inc.; and Verily Life Sciences. “ACCELERATING MEDICINES PARTNERSHIP and AMP are registered service marks of the U.S. Department of Health and Human Services.”

“Clinical data and biosamples used in preparation of this article were obtained from the (i) Michael J. Fox Foundation for Parkinson’s Research (MJFF) and National Institutes of Neurological Disorders and Stroke (NINDS) BioFIND study, (ii) Harvard Biomarkers Study (HBS), (iii) National Institute on Aging (NIA) International Lewy Body Dementia Genetics Consortium Genome Sequencing in Lewy Body Dementia Case-control Cohort (LBD), (iv) MJFF LRRK2 Cohort Consortium (LCC), (v) NINDS Parkinson’s Disease Biomarkers Program (PDBP), (vi) MJFF Parkinson’s Progression Markers Initiative (PPMI), and (vii) NINDS Study of Isradipine as a Disease-modifying Agent in Subjects With Early Parkinson Disease, Phase 3 (STEADY-PD3) and (viii) the NINDS Study of Urate Elevation in Parkinson’s Disease, Phase 3 (SURE-PD3).

“BioFIND is sponsored by The Michael J. Fox Foundation for Parkinson’s Research (MJFF) with support from the National Institute for Neurological Disorders and Stroke (NINDS). The BioFIND Investigators have not participated in reviewing the data analysis or content of the manuscript. For up-to-date information on the study, visit michaeljfox.org/news/biofind.”

“Genome sequence data for the Lewy body dementia case-control cohort were generated at the Intramural Research Program of the U.S. National Institutes of Health. The study was supported in part by the National Institute on Aging (program #: 1ZIAAG000935) and the National Institute of Neurological Disorders and Stroke (program #: 1ZIANS003154).”

“The Harvard Biomarker Study (HBS) is a collaboration of HBS investigators [full list of HBS investigators found at https://www.bwhparkinsoncenter.org/biobank/ and funded through philanthropy and NIH and Non-NIH funding sources. The HBS Investigators have not participated in reviewing the data analysis or content of the manuscript.”

“Data used in preparation of this article were obtained from The Michael J. Fox Foundation sponsored LRRK2 Cohort Consortium (LCC). The LCC Investigators have not participated in reviewing the data analysis or content of the manuscript. For up-to-date information on the study, visit https://www.michaeljfox.org/biospecimens).“

“PPMI is sponsored by The Michael J. Fox Foundation for Parkinson’s Research and supported by a consortium of scientific partners: 4D Pharma, AbbVie Inc., AcureX Therapeutics, Allergan, Amathus Therapeutics, Aligning Science Across Parkinson’s (ASAP), Avid Radiopharmaceuticals, Bial Biotech, Biogen, BioLegend, Bristol Myers Squibb, Calico Life Sciences LLC, Celgene Corporation, DaCapo Brainscience, Denali Therapeutics, The Edmond J. Safra Foundation, Eli Lilly and Company, GE Healthcare, GlaxoSmithKline, Golub Capital, Handl Therapeutics, Insitro, Janssen Pharmaceuticals, Lundbeck, Merck & Co., Inc., Meso Scale Diagnostics, LLC, Neurocrine Biosciences, Pfizer Inc., Piramal Imaging, Prevail Therapeutics, F. Hoffmann-La Roche Ltd and its affiliated company Genentech Inc., Sanofi Genzyme, Servier, Takeda Pharmaceutical Company, Teva Neuroscience, Inc., UCB, Vanqua Bio, Verily Life Sciences, Voyager Therapeutics, Inc., Yumanity Therapeutics, Inc.. The PPMI investigators have not participated in reviewing the data analysis or content of the manuscript. For up-to-date information on the study, visit www.ppmi-info.org.“

“The Parkinson’s Disease Biomarker Program (PDBP) consortium is supported by the National Institute of Neurological Disorders and Stroke (NINDS) at the National Institutes of Health. A full list of PDBP investigators can be found at https://pdbp.ninds.nih.gov/policy. The PDBP investigators have not participated in reviewing the data analysis or content of the manuscript.”

“The Study of Isradipine as a Disease-modifying Agent in Subjects With Early Parkinson Disease, Phase 3 (STEADY-PD3) is funded by the National Institute of Neurological Disorders and Stroke (NINDS) at the National Institutes of Health with support from The Michael J. Fox Foundation and the Parkinson Study Group. For additional study information, visit https://clinicaltrials.gov/ct2/show/study/NCT02168842. The STEADY-PD3 investigators have not participated in reviewing the data analysis or content of the manuscript.”

“The Study of Urate Elevation in Parkinson’s Disease, Phase 3 (SURE-PD3) is funded by the National Institute of Neurological Disorders and Stroke (NINDS) at the National Institutes of Health with support from The Michael J. Fox Foundation and the Parkinson Study Group. For additional study information, visit https://clinicaltrials.gov/ct2/show/NCT02642393. The SURE-PD3 investigators have not participated in reviewing the data analysis or content of the manuscript.”

“Data used in the preparation of this article were obtained from Global Parkinson’s Genetics Program (GP2). GP2 is funded by the Aligning Science Against Parkinson’s (ASAP) Initiative and implemented by The Michael J. Fox Foundation for Parkinson’s Research (https://www.gp2.org). For a complete list of GP2 members see http://www.gp2.org.“

## Notes

### Competing Interest Statement

The authors have declared no competing interest.

## References

1. Prinz, M., Jung, S. & Priller, J. Microglia biology: one century of evolving concepts. Cell 179, 292–311 (2019).

2. Kettenmann, H., Kirchhoff, F. & Verkhratsky, A. Microglia: new roles for the synaptic stripper. Neuron 77, 10–18 (2013).

3. Neumann, H., Kotter, M. R. & Franklin, R. J. M. Debris clearance by microglia: an essential link between degeneration and regeneration. Brain 132, 288–295 (2009).

4. Hu, X. et al. Microglial and macrophage polarization—new prospects for brain repair. Nat. Rev. Neurol. 11, 56–64 (2015).

5. Béchade, C., Cantaut-Belarif, Y. & Bessis, A. Microglial control of neuronal activity. Front. Cell. Neurosci. 7, 32 (2013).

6. Wolf, S. A., Boddeke, H. W. G. M. & Kettenmann, H. Microglia in Physiology and Disease. Annu. Rev. Physiol. 79, 619–643 (2017).

7. Hickman, S., Izzy, S., Sen, P., Morsett, L. & El Khoury, J. Microglia in neurodegeneration. Nat. Neurosci. 21, 1359–1369 (2018).

8. Hopp, S. C. et al. The role of microglia in processing and spreading of bioactive tau seeds in Alzheimer’s disease. J. Neuroinflammation 15, 1–15 (2018).

9. Scheiblich, H. et al. Microglia jointly degrade fibrillar alpha-synuclein cargo by distribution through tunneling nanotubes. Cell 184, 5089–5106 (2021).

10. Del Rio-hortega, P. El tercer elemento de los centros nerviosos. I. La microglia en estado normal II. Intervencion de la microglia en los procesos patologicos. HI. Naturaleza probable de la microglia. Boll Socieded Esp Biol 9, 69–120 (1919).

11. Lawson, L. J., Perry, V. H., Dri, P. & Gordon, S. Heterogeneity in the distribution and morphology of microglia in the normal adult mouse brain. Neuroscience 39, 151–170 (1990).

12. Villa, A., Vegeto, E., Poletti, A. & Maggi, A. Estrogens, neuroinflammation, and neurodegeneration. Endocr. Rev. 37, 372–402 (2016).

13. Matejuk, A. & Ransohoff, R. M. Crosstalk between astrocytes and microglia: an overview. Front. Immunol. 11, 1416 (2020).

14. Stratoulias, V., Venero, J. L., Tremblay, M. & Joseph, B. Microglial subtypes: diversity within the microglial community. EMBO J. 38, e101997 (2019).

15. Masuda, T., Sankowski, R., Staszewski, O. & Prinz, M. Microglia heterogeneity in the single-cell era. Cell Rep. 30, 1271–1281 (2020).

16. Schwarz, J. M., Sholar, P. W. & Bilbo, S. D. Sex differences in microglial colonization of the developing rat brain. J. Neurochem. 120, 948–963 (2012).

17. Weinhard, L. et al. Sexual dimorphism of microglia and synapses during mouse postnatal development. Dev. Neurobiol. 78, 618–626 (2018).

18. Crain, J. M., Nikodemova, M. & Watters, J. J. Microglia express distinct M1 and M2 phenotypic markers in the postnatal and adult central nervous system in male and female mice. J. Neurosci. Res. 91, 1143–1151 (2013).

19. Villa, A. et al. Sex-Specific Features of Microglia from Adult Mice. Cell Rep. 23, 3501–3511 (2018).

20. Hanamsagar, R. et al. Generation of a microglial developmental index in mice and in humans reveals a sex difference in maturation and immune reactivity. Glia 65, 1504–1520 (2017).

21. Vegeto, Elisabetta, Ciana, P. & Maggi, A. Estrogen and inflammation: hormone generous action spreads to the brain. Mol. Psychiatry 7, 236–238 (2002).

22. Gillies, G. E., Pienaar, I. S., Vohra, S. & Qamhawi, Z. Sex differences in Parkinson’s disease. Front. Neuroendocrinol. 35, 370–384 (2014).

23. Mazure, C. M. & Swendsen, J. Sex differences in Alzheimer’s disease and other dementias. Lancet. Neurol. 15, 451 (2016).

24. Beeson, P. B. Age and sex associations of 40 autoimmune diseases. Am. J. Med. 96, 457–462 (1994).

25. McCombe, P. A. & Henderson, R. D. Effects of gender in amyotrophic lateral sclerosis. Gend. Med. 7, 557–570 (2010).

26. Schapira, A. H. V. Glucocerebrosidase and Parkinson disease: recent advances. Mol. Cell. Neurosci. 66, 37–42 (2015).

27. Neumann, J. et al. Glucocerebrosidase mutations in clinical and pathologically proven Parkinson’s disease. Brain 132, 1783–1794 (2009).

28. Beutler, E. Gaucher disease: new molecular approaches to diagnosis and treatment. Science. 256, 794–799 (1992).

29. Neudorfer, O. et al. Occurrence of Parkinson’s syndrome in type 1 Gaucher disease. QJM An Int. J. Med. 89, 691–694 (1996).

30. Brunialti, E. et al. Inhibition of microglial β-glucocerebrosidase hampers the microglia-mediated antioxidant and protective response in neurons. J. Neuroinflammation 18, 1–18 (2021).

31. Garcia, J. A., Cardona, S. M. & Cardona, A. E. Analyses of microglia effector function using CX3CR1-GFP knock-in mice. in Microglia 307–317 (Springer, 2013).

32. Walter, T. et al. Visualization of image data from cells to organisms. Nat. Methods 7, S26–S41 (2010).

33. Zanier, E. R., Fumagalli, S., Perego, C., Pischiutta, F. & De Simoni, M.-G. Shape descriptors of the “never resting” microglia in three different acute brain injury models in mice. Intensive Care Med. Exp. 3, 0–18 (2015).

34. Elliot, E. J. & Muller, K. J. Long-term survival of glial segments during nerve regeneration in the leech. Brain Res. 218, 99–113 (1981).

35. Kloss, C. U. A., Bohatschek, M., Kreutzberg, G. W. & Raivich, G. Effect of lipopolysaccharide on the morphology and integrin immunoreactivity of ramified microglia in the mouse brain and in cell culture. Exp. Neurol. 168, 32–46 (2001).

36. Sheppard, O., Coleman, M. P. & Durrant, C. S. Lipopolysaccharide-induced neuroinflammation induces presynaptic disruption through a direct action on brain tissue involving microglia-derived interleukin 1 beta. J. Neuroinflammation 16, 1–13 (2019).

37. Villa, A., Rizzi, N., Vegeto, E., Ciana, P. & Maggi, A. Estrogen accelerates the resolution of inflammation in macrophagic cells. Sci. Rep. 5, 1–14 (2015).

38. Abd-El-Basset, E. & Fedoroff, S. Effect of bacterial wall lipopolysaccharide (LPS) on morphology, motility, and cytoskeletal organization of microglia in cultures. J. Neurosci. Res. 41, 222–237 (1995).

39. Nakamura, Y., Si, Q. S. & Kataoka, K. Lipopolysaccharide-induced microglial activation in culture: temporal profiles of morphological change and release of cytokines and nitric oxide. Neurosci. Res. 35, 95–100 (1999).

40. Masuda, T. et al. Spatial and temporal heterogeneity of mouse and human microglia at single-cell resolution. Nature 566, 388–392 (2019).

41. Pepe, G., Calderazzi, G., De Maglie, M., Villa, A. M. & Vegeto, E. Heterogeneous induction of microglia M2a phenotype by central administration of interleukin-4. J. Neuroinflammation 11, 1–14 (2014).

42. Kettenmann, H., Uwe Karsten, H., Mami, N. & Alexei, V. Physiology of microglia. Physiol. Rev. 91, 461–553 (2011).

43. Kuo, C. L. et al. In vivo inactivation of glycosidases by conduritol B epoxide and cyclophellitol as revealed by activity-based protein profiling. FEBS J. 286, 584–600 (2019).

44. Dermentzaki, G., Dimitriou, E., Xilouri, M., Michelakakis, H. & Stefanis, L. Loss of β-Glucocerebrosidase Activity Does Not Affect Alpha-Synuclein Levels or Lysosomal Function in Neuronal Cells. PLoS One 8, (2013).

45. Rocha, E. M. et al. Sustained Systemic Glucocerebrosidase Inhibition Induces Brain α-Synuclein Aggregation, Microglia and Complement C1q Activation in Mice. Antioxidants Redox Signal. 23, 550–564 (2015).

46. Vardi, A. et al. Delineating pathological pathways in a chemically induced mouse model of Gaucher disease. J. Pathol. 239, 496–509 (2016).

47. Rizzi, N., Rebecchi, M., Levandis, G., Ciana, P. & Maggi, A. Identification of novel loci for the generation of reporter mice. Nucleic Acids Res. 45, (2017).

48. Rizzi, N. et al. In vivo imaging of early signs of dopaminergic neuronal death in an animal model of Parkinson’s disease. Neurobiol Dis. 114, 74–84 (2018).

49. Tay, T. L., Savage, J. C., Hui, C. W., Bisht, K. & Tremblay, M. Microglia across the lifespan: from origin to function in brain development, plasticity and cognition. J. Physiol. 595, 1929–1945 (2017).

50. Wolf, S. A., Boddeke, H. & Kettenmann, H. Microglia in physiology and disease. Annu. Rev. Physiol. 79, 619–643 (2017).

51. Karperien, A. L., Jelinek, H. F. & Buchan, A. M. Box-counting analysis of microglia form in schizophrenia, Alzheimer’s disease and affective disorder. Fractals 16, 103–107 (2008).

52. Sierra, A., Paolicelli, R. C. & Kettenmann, H. Cien Años de Microglía: Milestones in a Century of Microglial Research. Trends Neurosci. 42, 778–792 (2019).

53. Zielasek, J. & Hartung, H.-P. Molecular mechanisms of microglial activation. Adv. Neuroimmunol. 6, 191–222 (1996).

54. Ewald, A. C. et al. Sex-and region-specific differences in the transcriptomes of rat microglia from the brainstem and cervical spinal cord. J. Pharmacol. Exp. Ther. 375, 210–222 (2020).

55. Guneykaya, D. et al. Transcriptional and translational differences of microglia from male and female brains. Cell Rep. 24, 2773–2783 (2018).

56. Han, J., Fan, Y., Zhou, K., Blomgren, K. & Harris, R. A. Uncovering sex differences of rodent microglia. J. Neuroinflammation 18, 1–11 (2021).

57. Villa, A., Della Torre, S. & Maggi, A. Sexual differentiation of microglia. Front. Neuroendocrinol. 52, 156–164 (2019).

58. Enquist, I. B. et al. Murine models of acute neuronopathic Gaucher disease. Proc. Natl. Acad. Sci. U. S. A. 104, 17483–17488 (2007).

59. Keatinge, M. et al. Glucocerebrosidase 1 deficient Danio rerio mirror key pathological aspects of human Gaucher disease and provide evidence of early microglial activation preceding alpha-synuclein-independent neuronal cell death. Hum. Mol. Genet. 24, 6640–6652 (2015).

60. Kaye, E. M., Ullman, M. D., Wilson, E. R. & Barranger, J. A. Type 2 and type 3 Gaucher disease: a morphological and biochemical study. Ann. Neurol. Off. J. Am. Neurol. Assoc. Child Neurol. Soc. 20, 223–230 (1986).

61. Norman, R. M., Urich, H. & Lloyd, O. C. The neuropathology of infantile Gaucher’s disease. J. Pathol. Bacteriol. 72, 121–131 (1956).

62. Hermida-Ameijeiras, Á., Méndez-Álvarez, E., Sánchez-Iglesias, S., Sanmartín-Suárez, C. & Soto-Otero, R. Autoxidation and MAO-mediated metabolism of dopamine as a potential cause of oxidative stress: role of ferrous and ferric ions. Neurochem. Int. 45, 103–116 (2004).

63. Corti, O., Lesage, S. & Brice, A. What genetics tells us about the causes and mechanisms of Parkinson’s disease. Physiol. Rev. (2011).

64. Li, Q., Jing, Y., Lun, P., Liu, X. & Sun, P. Association of gender and age at onset with glucocerebrosidase associated Parkinson’s disease: a systematic review and meta-analysis. Neurol. Sci. 42, 2261–2271 (2021).

65. Rosenbloom, B. et al. The incidence of Parkinsonism in patients with type 1 Gaucher disease: data from the ICGG Gaucher Registry. Blood Cells, Mol. Dis. 46, 95–102 (2011).

66. Simuni, T. et al. Clinical and dopamine transporter imaging characteristics of leucine rich repeat kinase 2 (LRRK2) and glucosylceramidase beta (GBA) Parkinson’s disease participants in the Parkinson’s progression markers initiative: a cross-sectional study. Mov. Disord. 35, 833–844 (2020).

67. Tan, E.-K. et al. Glucocerebrosidase mutations and risk of Parkinson disease in Chinese patients. Arch. Neurol. 64, 1056–1058 (2007).

68. Tan, Y.-L., Yuan, Y. & Tian, L. Microglial regional heterogeneity and its role in the brain. Mol. Psychiatry 25, 351–367 (2020).

69. Bradford, M. M. A rapid and sensitive method for the quantitation of microgram quantities of protein utilizing the principle of protein-dye binding. Anal. Biochem. 72, 248–254 (1976).

70. Toffoli, M. et al. Comprehensive short and long read sequencing analysis for the Gaucher and Parkinson’s disease-associated GBA gene. Commun. Biol. 5, 1–10 (2022).

