## Supplementary material for "Sex-specific microglial responses to glucocerebrosidase inhibition: relevance to GBA1-linked Parkinson disease": suppl table 1 and suppl fig 1


**This PDF file includes:**

Supplementary table 1

Supplementary figure 1

**Supplementary table 1**: cluster generated for each couple of parameters.

| COUPLE OF PARAMETERS | | CLUSTERS GENERATED |
|---|---|---|
| Area-Motility | Area↓ Motility↓ | Small & Static |
| | Area↓ Motility ↑ | Small & Motile |
| | Area↑ Motility ↓ | Big & Static |
| | Area↑ Motility ↑ | Big & Motile |
| CV% Solidity-Motility | CV% Solidity↓ Motility↓ | Steady & Static |
| | CV% Solidity↓ Motility↑ | Steady & Motile |
| | CV% Solidity↑ Motility↓ | Variable & Static |
| | CV% Solidity↑ Motility↑ | Variable & Motile |
| CV Area - Solidity | CV Area↓ - Solidity↓ | Inactive & Complex |
| | CV Area ↓- Solidity↑ | Inactive & Simple |
| | CV Area↑ - Solidity↓ | Contractile & Complex |
| | CV Area↑ - Solidity↑ | Contractile & Simple |
| Solidity - Area | Solidity↓ Area↓ | Complex & Small |
| | Solidity ↓ Area↑ | Complex & Big |
| | Solidity ↑ Area↓ | Simple & Small |
| | Solidity↑ Area↑ | Simple & Big |
| Solidity - Motility | Solidity↓ Motility↓ | Complex & Static |
| | Solidity↓ Motility↑ | Complex & Motile |
| | Solidity↑ Motility↓ | Simple & Static |
| | Solidity ↑ Motility↑ | Simple & Motile |
| Rotation - Motility | Rotation↓ Motility↑ | Stationary & Static |
| | Rotation↓ Motility↓ | Stationary & Motile |
| | Rotation ↑ Motility↑ | Rotant & Static |
| | Rotation ↑ Motility↓ | Rotant & Motile |

↓ values under the median; ↑ values over the median

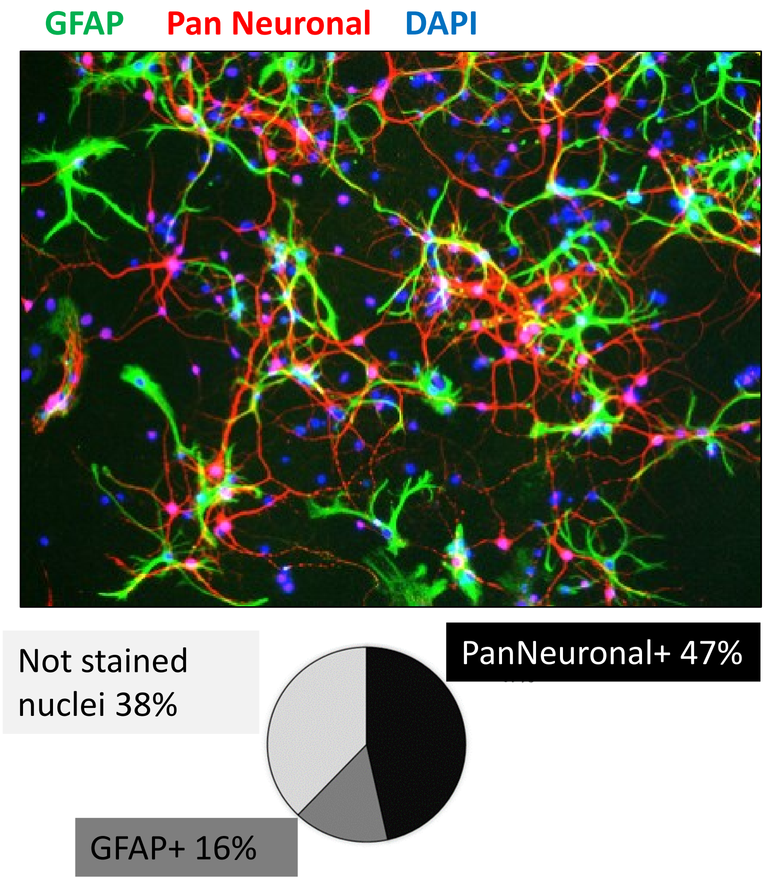


**Supplementary figure 1:** primary cortical cells layer composition to supports microglial function. Top) representative fluorescent image of the cortical neuronal cultures. green= GFAP marker, red=pan-neuronal marker, blue=DAPI. Bottom) the percentage of cell composition is reported with respect to the total nucleus counted.
